# Assessing by modeling the consequences of increased recombination in genomic selection of *Oryza sativa* and *Brassica rapa*

**DOI:** 10.1101/704544

**Authors:** E. Tourrette, R. Bernardo, M. Falque, O. Martin

**Author notes:** corresponding author: Olivier C. Martin, GQE – Le Moulon, INRA, Univ. Paris-Sud, CNRS, AgroParisTech, Université Paris-Saclay, Gif-sur-Yvette, France.

## Abstract

Recombination generates genetic diversity but the number of crossovers per meiosis is limited in most species. Previous studies showed that increasing recombination can enhance response to selection. However, such studies did not assume a specific method of modifying recombination. Our objective was to test whether two methods used to increase recombination in plants could increase the genetic gain in a population undergoing genomic selection. The first method, in *Oryza sativa,* used a mutant of anti-crossover genes to increase global recombination without affecting the recombination landscape. The second one uses the ploidy level of a cross between *Brassica rapa* and *Brassica napus* to increase the recombination particularly in pericentromeric regions. These recombination landscapes were used to model recombination while quantitative trait loci positions were based on the actual gene distribution. We simulated selection programs with initially a cross between two inbred lines, for two species. Increased recombination enhanced the response to selection. The amount of enhancement in the cumulative gain largely depended on the species and the number of quantitative trait loci (2, 10, 20, 50, 200 or 1000 per chromosome). Genetic gains were increased up to 30% after 20 generations. Furthermore, modifying the recombination landscape was the most effective: the gain was larger by 25% with the first method and 33% with the second one in *B. rapa*, and 15% compared to 11% in *O. sativa*. Thus, increased recombination enhances the genetic gain in genomic selection for long-term selection programs, with visible effects after four to five generations.

## INTRODUCTION

Recombination is one of the processes generating genetic diversity, creating new *allelic combinations* through sexual reproduction. Hence, recombination is a cornerstone for obtaining genetic progress during selection. Aside from providing genetic diversity driving genetic progress, recombination is key to a number of practical applications. For instance, recombination intensity controls the resolution of genetic mapping and QTL analyses (**Lander and Botstein 1989, van Ooijen 1992**) and it is particularly important for creating specific mapping populations such as Recombinant Inbred Lines (RILs) (**Bailey 1971** for the development of RILs in mouse, **Lister and Dean 1993** for the development of RILs for mapping in *A. thaliana*).

Theoretically, it has been shown that recombination is required for the efficiency of selection, by breaking the linkage between QTLs (**Hill and Robertson 1966, Felsenstein 1974**). Indeed, when two QTLs are linked, they tend to be inherited together. If the alleles of these QTLs have opposite effects on the traits of interest, that can impede the efficiency of selection. Such negative linkage disequilibrium can be generated in finite populations due to selection (Bulmer effect, **Bulmer 1971**) or genetic drift (**Barton 2009**).

A consequence of increased recombination in selection is the faster accumulation of advantageous mutations in evolving populations. In the limit, for instance, of a population without recombination, it is difficult to get rid of deleterious mutations even in the presence of selection (**Muller 1964, Felsenstein 1974**). The relevance of modifying recombination has also been seen in some experimental studies of the rate of response to selection (**McPhee and Robertson 1970, Zeyl and Bell 1997,** see **Burt 2000** for a review**, Goddard *et al*. 2005, Lovell *et al*. 2014**). These studies showed what would happen if one were to suppress recombination.

Inversely, based on such theoretical considerations, could one have an increase in the efficiency of selection by *increasing* recombination rate? This question is all the more interesting when considering that recombination rate is limited in most species, with there almost never being more than 3 crossovers (COs) per meiosis per chromosome pair regardless of the chromosome physical size (**Mercier *et al*. 2015, Fernandes *et al*. 2018**). Two studies using simulations looked at the effects of increasing recombination on genetic gain during selection programs.

**McClosky and Tanksley (2013)** simulated a plant breeding experiment with a biparental initial population. They first looked at what could be gained by having all loci segregating independently, hereafter referred to as “free recombination”, compared to normal recombination: they found that the average gain was 11%. They then thought about ways to either accumulate recombination events or increase recombination. To accumulate recombination events, they did more rounds of random mating before selection. To increase recombination, they selected higher recombinant individuals in parallel to selecting on their trait of interest. In this way, they found they could attain at most half of the genetic progress that was obtained in the case of free recombination and they concluded that the most effective technique was to perform this simultaneous double selection on both the trait of interest and recombination rate. **Battagin *et al*. (2016)** simulated a livestock breeding experiment in which there were two successive phases, with first the production of a historical population to get a genetic structure similar to that of an actual population used for breeding, and then a breeding program based on that population. They looked at different values for the recombination rates by increasing the genetic lengths of the chromosomes. For a 2-fold increase of the genetic lengths, they found that the gain was 12.5% higher when considering a program covering 40 generations, for a 10-fold increase the gain was 28.7% higher while for a 20-fold increase the gain was 33.4% higher. Hence, to get a high increase of the gain, they had to increase the recombination rate by 10 to 20-fold. The study by **Battagin *et al*.** found a higher increase than the one by **McClosky and Tanksley**, which could be explained by the population structure (initial diversity) or by the nature of the selection program (in livestock, distinction between males and females, selection on the males only). **Gonen *et al. (*2017)**, while not interested in increased recombination rate, looked at shifting recombination hotspots to have access to more allelic variations. They observed an increase in the genetic gain when shifting QTLs from outside to inside hotspots. That led to a similar conclusion as the previous studies, namely that it may be beneficial to increase recombination rates everywhere in the genome.

Since these studies were performed, different methods of modifying recombination rate along the genome have been developed (see **Blary and Jenczewski 2018** for a review of the different methods). By modifying recombination, we mean either globally increasing the recombination rate (more relevant to the first two simulation studies) or modifying the recombination landscape (more relevant to the third simulation study mentioned above). A first method, the so-called HyperRec technology, is based on suppressing genes that have an anti-crossover role. To date, three associated pathways have been used, corresponding to the knock-out of three genes: *FANCM* (**Crismani *et al*. 2012**), *RECQ4* (**Séguéla-Arnaud *et al*. 2015**) and *FIGL1* (**Girard *et al*. 2015**). When these genes are knocked out, one sees major increases in the number of crossovers, though without changing the recombination landscape, be it in *A. thaliana* (**Fernandes *et al*. 2018**), pea, rice or tomato (**Mieulet *et al*. 2018)**. For instance, the double mutant *recq4 fancm* showed 7.8-fold increase in the number of COs compared to the wild type in *thaliana.* Another situation leading to increased recombination has been discovered in polyploids. Specifically, it has been found that the ploidy levels in *Brassica* species have a strong influence both on global recombination rates and on recombination landscapes. For instance allotriploid hybrids AAC (single copy of the C genome), obtained by crossing *B. rapa* (diploid AA) with *B. napus* (allotetraploid AACC), produced crossovers in the A genome at rates about 3 times those arising in *B. rapa* diploids. More interestingly, it led to a tremendous increase in the recombination rate within the pericentromeric regions where neither *B. rapa* nor *B. napus* recombine (**Leflon *et al*. 2010, Suay *et al*. 2014, Pelé *et al*. 2017**).

Other methods for increasing recombination rates include genome editing to target COs to specific genomic sites (see **Hayut *et al*. 2017** for an application of genome editing in tomato, **Bernardo 2017**, **Ru and Bernardo 2018**, **Brandariz and Bernardo 2019**, for assessing the potential use of targeted recombination in plant breeding via simulations) or over-expressing pro-crossover genes such as HE10 (**Ziolkowski *et al*. 2017, Serra *et al*. 2018**). It is also known that modifications of the growth environment can affect recombination rates. For example, recombination is sensitive to the temperature (see **Phillips *et al*. 2015** and **Lloyd *et al*. 2018** for examples in barley and *A. thaliana*, **Modlizewski and Copenhaver 2017** for a review on different stresses affecting recombination). There is also some variability across individuals within a given species, for instance associated with the sex context of meiosis (**Giraut *et al*. 2011, Phillips *et al*. 2015** for examples in *A. thaliana* and barley respectively) or from genetic differences between the individuals as revealed by intra-specific variability of recombination rates and landscapes (**Salomé *et al*. 2011, Bauer *et al*. 2013**, for examples in *A. thaliana* and maize respectively).

In this study, we investigate to what extent is it interesting for breeding programs to modify recombination, considering the first two experimental methods mentioned above, that is (i) using mutants of anti-crossover genes and (ii) changing ploidy levels in *Brassicae*. For that we shall work with the actual recombination profiles experimentally observed with each of these two methods. For our simulations, we will expand on the approach used by **McClosky and Tanksley (2013)**, using a biparental population and parameters in a range similar to the ones they used. We will investigate the effects of a number of parameters, in particular the number of QTLs (2, 10, 20, 50, 200 and 1000 per chromosome), the presence of coupling or repulsion between QTLs, crossover interference, and the level of trait heritability (h^2^ = 0.2, 0.5, 0.8 and 1). Finally, instead of using only the phenotype for selection, we also test the consequences of modified recombination when using genomic selection.

## MATERIALS AND METHODS

We investigated the effects of modifying recombination on genetic gain in selection programs by mathematical and computer modeling. All our codes were written in the programming language R, both for producing individual-based simulations of forward-in-time selection programs and for all the associated analyses.

### HyperRec (O. sativa) and boosted (B. rapa) recombination landscapes

The normal and modified recombination landscapes used in our simulations were taken from the published results in *Brassica rapa* (turnip, **Pelé *et al*. 2017**) and *Oryza sativa* (rice, **Mieulet *et al*. 2018**). In the case of *B. rapa*, the modified recombination landscape has an increased recombination rate in the pericentromeric regions which are otherwise poor in crossovers. For *O. sativa*, the recombination rate is globally increased but without significant changes in the recombination profile, the pericentromeric regions still being poor in crossovers.

For *B. rapa*, the modified recombination will be called **boosted** recombination and for *O. sativa* it will be called **HyperRec** (HR) recombination. As *O. sativa* and *B. rapa* have different genetic parameters (number of chromosomes, genetic lengths), expected HR and boosted recombination landscapes were also simulated for *B. rapa* and *O. sativa* respectively (see the paragraph on the construction of the landscapes). The recombination profiles in these different situations are presented in Fig1 and Fig2.

**Figure 1:**
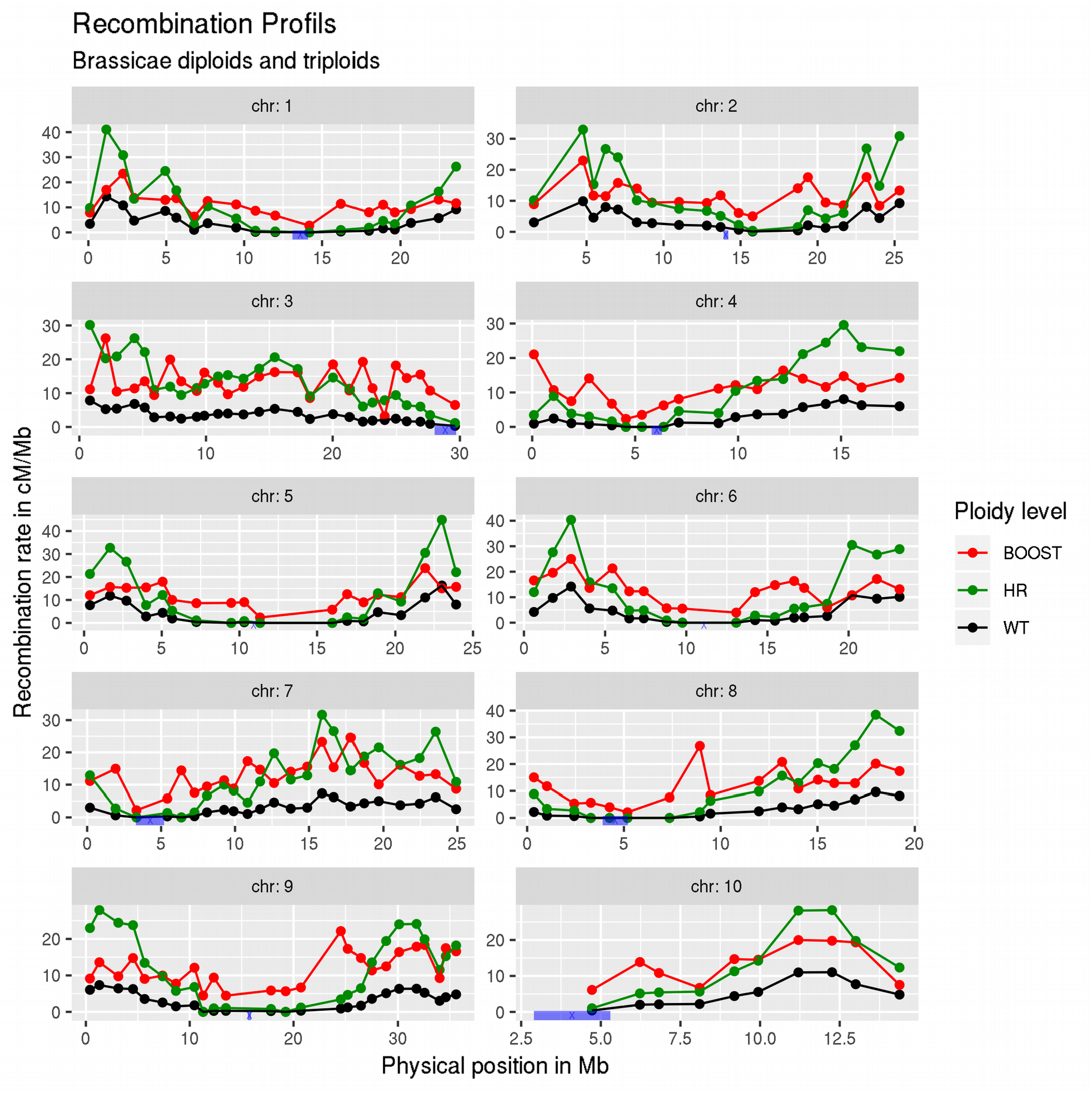
Recombination landscape for each chromosome of *B. rapa* for normal (black), HR (green) and boosted (red) recombination. The HR landscape was defined so that its profile was the same as for normal recombination, while the normal and boosted landscapes were from **Pelé *et al*. (2017)**. In blue, *B. rapa* centromere position from **Mason *et al*. (2016)**.

**Figure 2:**
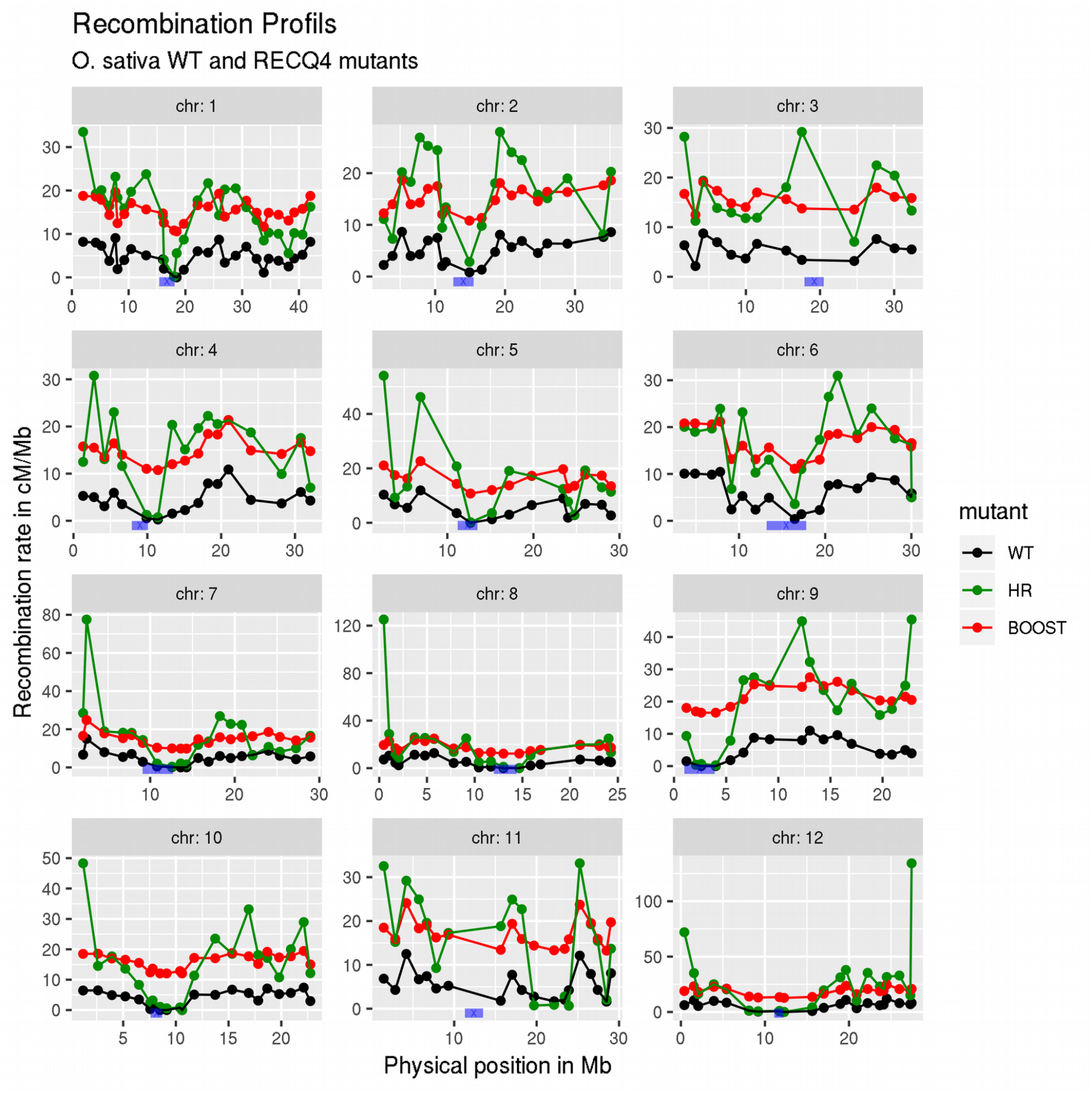
Recombination landscape for each chromosome of *O. sativa* for normal (black), HR (green) and boosted (red) recombination. The boosted recombination landscape was defined *via* an additive increase everywhere while the normal and HR landscapes were from **Mieulet *et al*. (2018)**. In blue, the centromere position taken from **Mizuno *et al*., (2018)**.

### QTLs effects

For each chromosome, we allowed for 2, 10, 20, 50, 200, or 1000 loci carrying biallelic QTLs affecting the trait considered. We considered these different cases for the numbers of QTLs to test different genetic architectures underlying our trait.

The amplitude of each QTL effect was drawn from a gamma distribution; our choice for the shape parameter k=0.4 leads to an L-shaped distribution so that most of the QTLs had a weak effect (**Hayes and Goddard 2001, Technow *et al*. 2012**). The gamma distribution’s scale parameter θ was chosen such that the variance of this QTL effect (equal to kθ^2^) was 1 and thus θ = 1.58. As the population is biallelic, each allelic effect was equal to half the QTLs’ amplitude. A sign was then randomly assigned for one allelic effect and the opposite sign was taken for the other one. The distributions of absolute allelic effects for 2, 10, 20, 50 and 200 QTLs per chromosome are shown in **FigS1.**

### Allowing for coupling or repulsion between QTLs

Recombination should benefit selection by breaking the linkage between the QTLs with an opposite allelic effect on the trait (repulsion case). On the contrary, recombination is expected to be detrimental to selection by breaking the positive correlations between two QTLs having both positive or negative effects (coupling case). Hence, we also looked at the effect of coupling or repulsion on the efficiency of boosted recombination. To model the coupling (respectively repulsion), we introduced a probability of changing (respectively keeping) the same sign as the previous QTL when creating each individual. This probability *P* is equal to 1/2(1 - *exp(-Δx/L))*, *Δx* being the genetic distance between the two QTLs and *L* being the characteristic genetic length of the coupling (respectively repulsion). The larger *L*, the further one will maintain significant correlations between allelic signs. Under coupling, *P* is the probability that the sign will *change* between adjacent QTLs; under repulsion, *P* is the probability that the sign will stay the same. When *L* tends to infinity or *Δx* tends to 0, *P = 0*. On the other hand, when *L* tends to 0 or *Δx* tends to infinity, *P = ½* so that there is no coupling nor repulsion between adjacent QTLs and the sign of an allele will be assigned independently of the sign of the previous QTL. In our situation, we choose a characteristic length *L = 5 cM*, which results in moderate coupling/repulsion. Such levels are undone rather quickly under boosted recombination.

### QTL positions

The positions of the QTLs were drawn independently with a density proportional to the gene density of each species (taking gene distributions without transposable elements; **FigS2** for the gene distribution of *B. rapa* and **FigS3** for *O. sativa*). Gene physical positions were downloaded from public databases (for *O. sativa*: https://rapdb.dna.affrc.go.jp/download/irgsp1.html, file “Gene structure information in GFF format” for the computationally predicted genes; and for *B. rapa*: http://brassicadb.org/brad/datasets/pub/BrassicaceaeGenome/Brassica_rapa/V3.0/). QTL physical positions were thus *sampled* at random among the gene physical positions, for each replicate of our simulations. As only genetic positions in centiMorgans were used during simulations, we had to convert the QTL physical positions into their genetic positions by using the genetic map for each situation (normal, HR, and boosted recombinations for both *O. sativa* and *B. rapa*).

### Construction of the HyperRec recombination landscape in B. rapa and the boosted recombination landscape in O. sativa

The genetic maps of *O. sativa* for the wild type (WT, with normal recombination) and *Osrecq4* mutant (HR recombination) were set using marker genetic and physical positions given as supplementary material in **Mieulet *et al*. (2018)**. The ones for *B. rapa* A_n_A_r_’ (female) diploid (normal recombination) and A_n_A_r_’C_n_ allotriploid (boosted recombination) were set using the marker physical positions taken from **Pelé *et al*. (2017)** and the genetic positions kindly communicated by Pelé.

Since the experimental landscapes of boosted recombination in *O. sativa* and HR recombination in *rapa* have not yet been measured, they were simulated. The expected boosted recombination landscape was defined by *adding* a constant to the recombination rate (in cM/Mb) of the normal recombination landscape (same increase of recombination everywhere along the chromosome). This constant was (GL_HR_ – GL_NR_)/PL, with GL_HR_ and GL_NR_ being the genetic lengths in cM under HR and normal recombination, and PL being the physical length of the chromosome in Mb. The HR recombination landscape was simulated by *multiplying* the normal recombination rate by a constant, GL_BR_/GL_NR_, with GL_BR_ and GL_NR_ being the genetic lengths under boosted and normal recombination (thus the increase of recombination is proportional to the initial recombination). Both of these constants were chosen to obtain the same total increase of recombination for HR and boosted recombination compared to normal recombination.

Given that the landscapes are specified experimentally by markers, it is necessary to interpolate between them to be able to simulate crossovers at arbitrary positions; for that, the relationships between genetic and physical positions were modeled using splines, from which QTL genetic positions were obtained. Gene physical positions used for sampling were restricted to the physical range of the genetic map as we do not have the relation between genetic and physical positions outside of this mapped region.

We took the same QTL *physical* positions and effects for normal and modified recombination, only changing the simulated QTL *genetic* positions based on the different recombination profiles (our simulations operate in genetic space).

The genetic lengths used in the different situations were given by the genetic maps and are shown in Table1.

**Table1:**
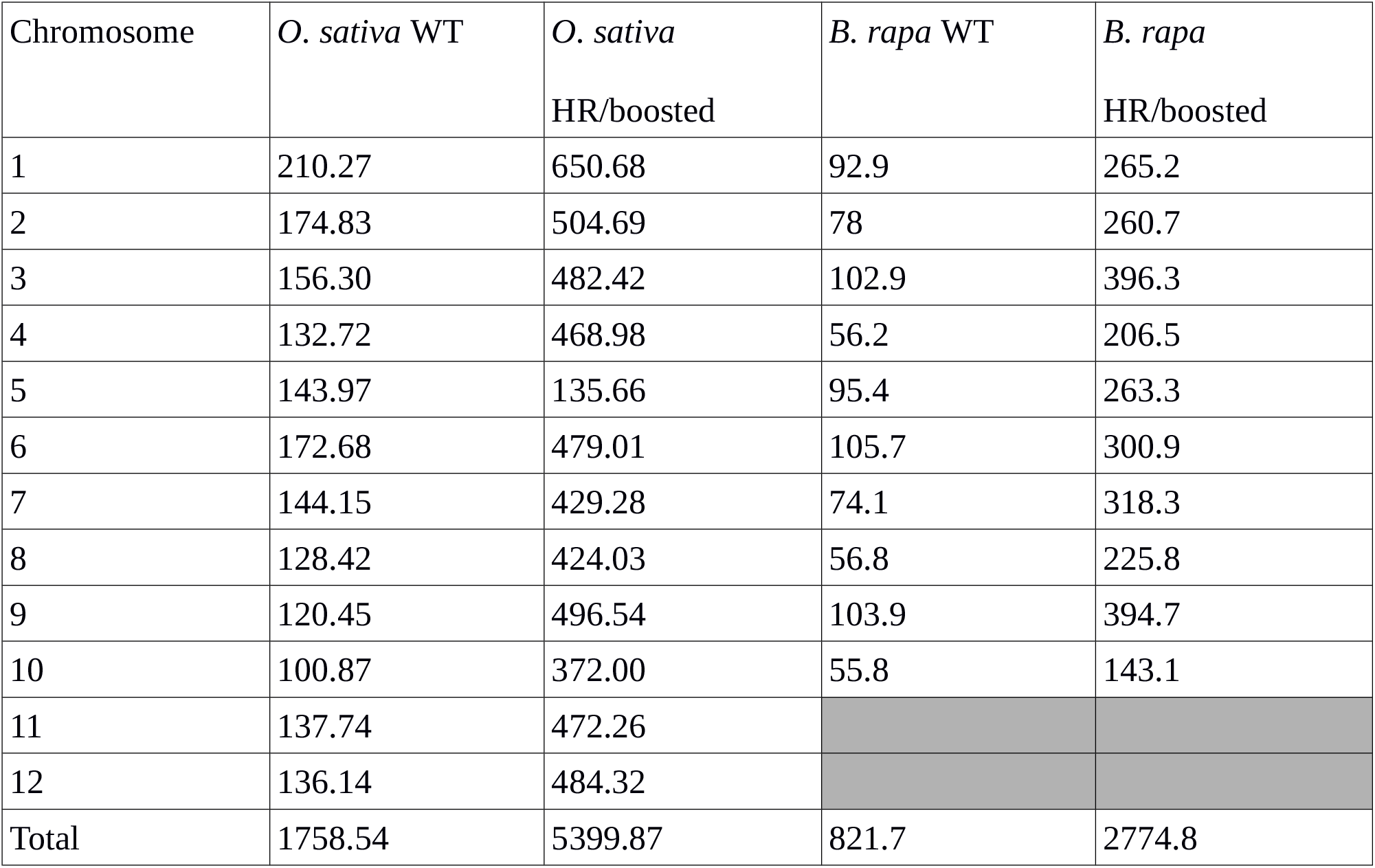
Genetic lengths, in centiMorgans, for the twelve chromosomes of *O. sativa* and the ten chromosomes of *B. rapa* under normal and increased recombination (HR and boosted). The genetic lengths for HR and boosted recombination were taken as identical, as we wanted the same global increase of recombination to be able to investigate specific effects of changing the recombination *landscape*.

| Chromosome | *O. sativa* WT | *O. sativa*<br>HR/boosted | *B. rapa* WT | *B. rapa*<br>HR/boosted |
| --- | --- | --- | --- | --- |
| 1 | 210.27 | 650.68 | 92.9 | 265.2 |
| 2 | 174.83 | 504.69 | 78 | 260.7 |
| 3 | 156.30 | 482.42 | 102.9 | 396.3 |
| 4 | 132.72 | 468.98 | 56.2 | 206.5 |
| 5 | 143.97 | 135.66 | 95.4 | 263.3 |
| 6 | 172.68 | 479.01 | 105.7 | 300.9 |
| 7 | 144.15 | 429.28 | 74.1 | 318.3 |
| 8 | 128.42 | 424.03 | 56.8 | 225.8 |
| 9 | 120.45 | 496.54 | 103.9 | 394.7 |
| 10 | 100.87 | 372.00 | 55.8 | 143.1 |
| 11 | 137.74 | 472.26 |  |  |
| 12 | 136.14 | 484.32 |  |  |
| Total | 1758.54 | 5399.87 | 821.7 | 2774.8 |

### Simulating crossover formation

Meiotic recombination was simulated by generating crossovers in genetic space. The number of crossovers was drawn from a Poisson distribution, as required by the no-interference model of Haldane (**McPeek and Speed, 1995**). The genetic distance between two adjacent crossovers was drawn from an exponential distribution, with an average of 1 crossover per 100 cM.

We also performed a study including interference which was modeled following the single-pathway (considers only interfering crossovers) Gamma model (**McPeek and Speed, 1995**), using a gamma distribution for the distance between adjacent crossovers. Hence, using this model with interference, it is possible to modify the “repulsion” of crossovers generated within a single meiosis, the strength of which depends on the *nu* parameter of the distribution. When this parameter is equal to 1, it is equivalent to the model of Haldane without interference. The values of the *nu* parameter for each chromosome in the diploid and in the allotriploid hybrid of *B. rapa* (normal and boosted recombination) were taken from **Pelé *et al*. 2017**. The values are shown in Table2. In **Pelé *et al*. (2017)**, interference was estimated using two-pathways instead of the single-pathway. Contrary to the single-pathway, both interfering and non-interfering crossovers are taken into account for the estimation of the *nu* parameter by those authors. As a result, by using the interference parameters of the two-pathway model in our single pathway approach, we probably overestimated the effect of interference (*nu* parameters higher than they should be).

**Table2:** Values of the *nu* parameters of the gamma distribution controlling the level of interference for the diploid (normal recombination) and allotriploid hybrids (boosted recombination) of *B. rapa* (taken from **Pelé et al. 2017).** These values are for the two-pathways gamma model (superposition of interfering and non-interfering crossovers) while we used the single pathway model. Hence these values overestimate the interference strength, so they can be considered as giving upper limits for the effects of the interference.

| Chromosome | Diploid AA | Allotriploid AAC |
| --- | --- | --- |
| A01 | 6.97 | 1.887 |
| A02 | 6.139 | 1.479 |
| A03 | 7.207 | 2.392 |
| A04 | 18.576 | 1.946 |
| A05 | 12.934 | 2.263 |
| A06 | 8.814 | 3.084 |
| A07 | 13.001 | 1.64 |
| A08 | 12.185 | 2.036 |
| A09 | 4.895 | 2.418 |
| A10 | 63.778 | 2.362 |

### Neutral markers

We also introduced 1000 biallelic neutral markers per chromosome (having no effect on the trait and regularly spaced on the genetic map). They were taken to be equally spaced, the genetic distance between adjacent markers being given by the ratio between the chromosome’s genetic length and the number of neutral markers minus 1. The number of these markers was kept constant while the genetic lengths changed according to the modified or not recombination landscapes.

### The quantitative genetics framework and selection schemes

For each individual we denote its genotypic value by *G*. As only additivity was considered, *G* was equal to the allelic effects present in the considered individual, summed over all QTLs and thus over all pairs of homologous chromosomes.

The initial population, bi-parental, was generated from two homozygous lines P1 and P2 (generation g = 0), with P1 having one of the two alleles for each QTL and P2 the other one. P1 and P2 were crossed to get an F1 individual (generation g = 1).

Two different selection schemes were tested: a “Fn selection scheme” and a “DH selection scheme”. In the Fn selection scheme, the F1 was selfed to produce a population of 250 F2 individuals (g = 2). From this F2 generation, each further generation was obtained *via* (i) a step of “truncation selection” to keep the “best” individuals within the population based on their breeding value, and (ii) a step of random mating between these selected individuals to produce the offspring that form the individuals of the next generation. We simulated this selection program over a total of g = 20 generations. The populations had a constant size of 250 individuals and the selection pressure was 2%, meaning that only the 5 best individuals were kept to create the next generation.

In the DH selection scheme, the F1 was used to get 250 doubled haploids (DH, g = 2), among which the 5 best individuals were then selected and crossed to get a new heterozygous population of the same size. Each one of these individuals was then used to produce one DH, and among these DHs the best ones were again selected and crossed for the next generation. These steps, producing respectively homozygous and heterozygous populations, were alternated during 20 generations, corresponding to 10 cycles of selection. The selection intensity was the same as in the Fn selection scheme, *i.e.*, 5 individuals among the 250 and the population size of 250 individuals was also constant.

One cycle of the Fn selection scheme is shown in Fig3 while one cycle of the DH selection scheme is shown in Fig4. As we performed up to 20 generations of selection, we also considered the selection of 10 and 20 individuals among the 250 (selection intensity of 4% and 8% respectively) to include more realistic situations.

**Figure 3:**
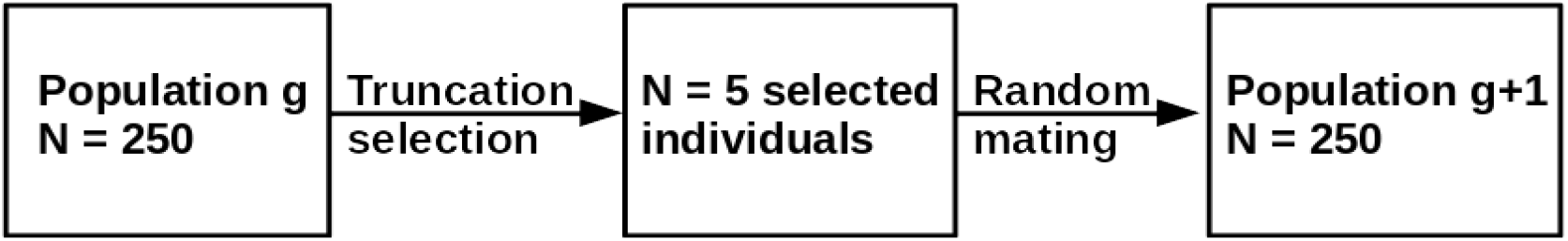
Schematic representation of one cycle of selection in the Fn selection scheme

**Figure 4:**
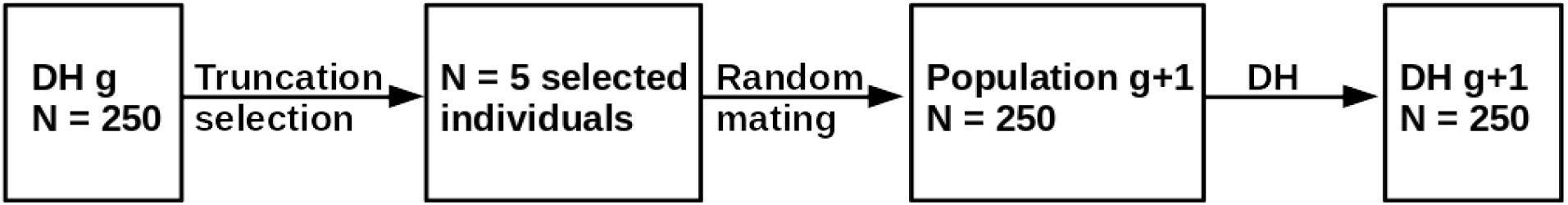
Schematic representation of one cycle of selection in the DH selection scheme

We considered two selection criteria where the “breeding value” corresponded either to the phenotypic value P (in the case of phenotypic selection) or to the genomic estimated breeding value GEBV (in the case of genomic selection).

For phenotypic selection, P was calculated as the genotypic value G plus an environmental noise E. E was taken from a normal distribution with null mean and a variance V_E_. To set V_E_, we first estimated the genetic variance V_G_ in the F2 population, and calculated V_E_ to get the desired value of the heritability defined as h^2^ = V_G_/V_P_ = V_G_/[V_G_+V_E_] (V_P_ being the phenotypic variance). Four different values of h^2^ were tested: 1, 0.8, 0.5 and 0.2. Note that h^2^ is defined based on the F2 population, the later populations having different variances will also have different values of h^2^.

In the case of genomic selection, GEBV was calculated using the estimated marker effects and the population mean. Both were estimated in the F2 population and then every fourth generation (training populations) with ridge regression best linear unbiased prediction (RR-BLUP) using the rrBLUP R package (**Endelman 2011**). This corresponds to the mixed model Y = Xβ + Zu + e, where Y is the vector (Nx1) of phenotypic values calculated as for phenotypic selection, X a vector (Nx1) of 1s, Z the genotype matrix (NxN_marker_) and e the vector of residuals. The genotype matrix was coded as follows: Z_ij_ = −1 if the marker j of individual i is homozygous for the allele of parent 1, Z_ij_ = 1 if it is homozygous for the allele of parent 2, and Z_ij_ = 0 if it is heterozygous. The population mean β and the vector of marker effects u were then estimated and used in the next generations to calculate GEBV = β + Z_gu_, where Z_g_ was the genotype matrix of the generation g, coded as previously described.

An important consideration in genomic selection is the frequency with which one re-calibrates the estimated marker effects along generations of selection. In this work we considered three such cases: a single calibration arising in the F2 generation, calibrations every fourth generation and calibrations every second generation.

### Parameter choices

Clearly there are many parameters to set in any framework such as ours. In the section Results, unless otherwise stated, the parameters used are those of the Fn selection scheme, under genomic selection with the marker effects estimated every fourth generation. Furthermore, we use 200 QTLs per chromosome and a trait heritability of 0.5, and no crossover interference. The selection intensity is set to 2% and we impose no coupling or repulsion. As justified in the Results section, the analyses will mainly focus on the species *B. rapa*. To understand the role of the different parameters in our framework, we consider them one at a time. Specifically, starting with the default values just given above, we see what happens when one changes one of our choices, such as the number of QTLs or the frequency of calibration of the genomic selection model.

### Using replicates in the simulations and statistical analyses

For each generation, the genetic gain was calculated as [G_g_-G_g_ = 2] where G denotes the population average of the genotypic values. We also calculated at each generation the genetic variance of the population as the variance of the vector of genotypic values. The *efficiency* of having modified recombination was calculated as the ratio of genetic gains: G_modified recombination_/ G_normal recombination_. To assess the stochastic variability of the process, we used 1000 replicates for each study (choice of selection program and parameters) and calculated their genetic gain, genetic variance and relative gain. Confidence intervals were then calculated via the standard error on the mean values of these quantities, 1.96*σ/√Nreplicates.

### Data Availability Statement

The scripts for the simulations are available as a R package on https://sourcesup.renater.fr/frs/download.php/latestfile/2217/CAREB_R.tar.gz The supplementary figures S1 to S10 have been deposited on figshare.

## RESULTS

We hypothesize that increased recombination can be beneficial for selection by (1) breaking linkage between QTLs and (2) providing access to genetic diversity by introducing recombination into cold regions. To test the first hypothesis, we studied different numbers of QTLs per chromosome: the more QTLs per chromosome there are, the tighter the linkage. Hence, as the number of QTLs per chromosome increases, the effect of increased recombination should also increase. Note that we are more interested in the number of QTLs per chromosome than in the total number of QTLs genome-wide: indeed, recombination arises within and across chromosomes but both HR and boosting act only on the former. To test the second hypothesis, we compared the increase of recombination by HR and boosted recombination, where the first method does not change significantly the recombination landscape (no crossovers in the cold regions in both normal and HR) while the second one changes the recombination landscape and introduces crossovers into the cold regions. Our modeling and simulational approaches are explained in the Methods section; a set of reference parameters are used (e.g., population size, number of QTLs, heritability, level of selection, etc) to illustrate the effects of increased recombination but of course we also explore the roles of these different parameters, showing in particular that these effects are almost always robust to changing the values of the parameters.

### Increased recombination leads to higher genetic gain

We find that increasing recombination slows down both the loss of genetic diversity and the decrease of genetic variance (Fig5). This results in an increase of the genetic gain and a delay in the fixation of the QTL alleles, the plateau for the genetic gain (due to fixation) being reached later (Fig6).

**Figure 5:**
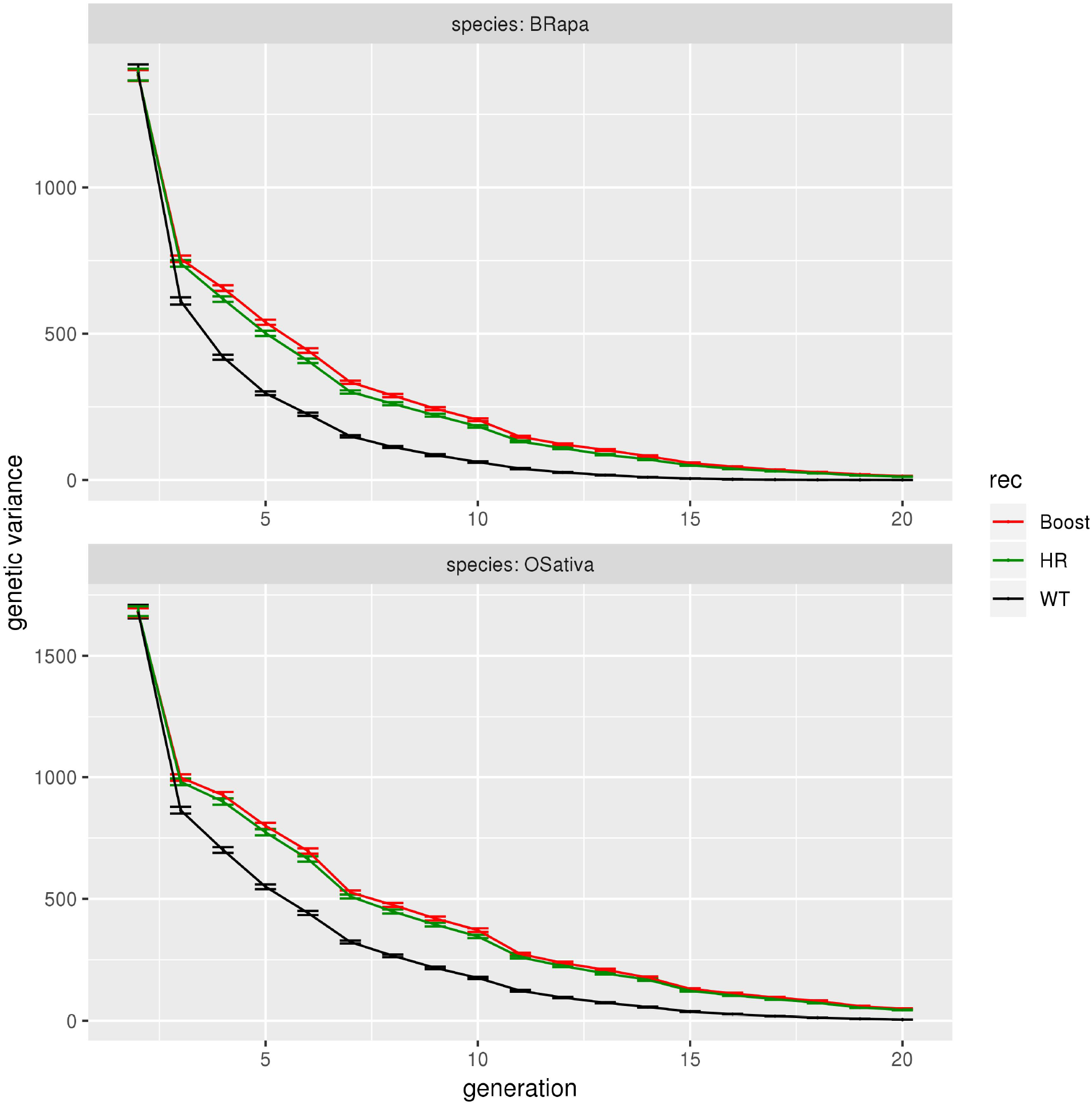
Genetic variance as a function of generation, for *B. rapa* and *O. sativa* under normal (black), HR (green) and boosted (red) recombination for genomic selection with the marker effects estimated every fourth generation, a heritability of 0.5, selection on heterozygotes with an intensity of 2%, 200 QTLs per chromosome, crossovers formed without interference, no coupling nor repulsion. The first generation shown is the F2 generation (g = 2). The error bars represent 95% confidence interval on the mean.

**Figure 6:**
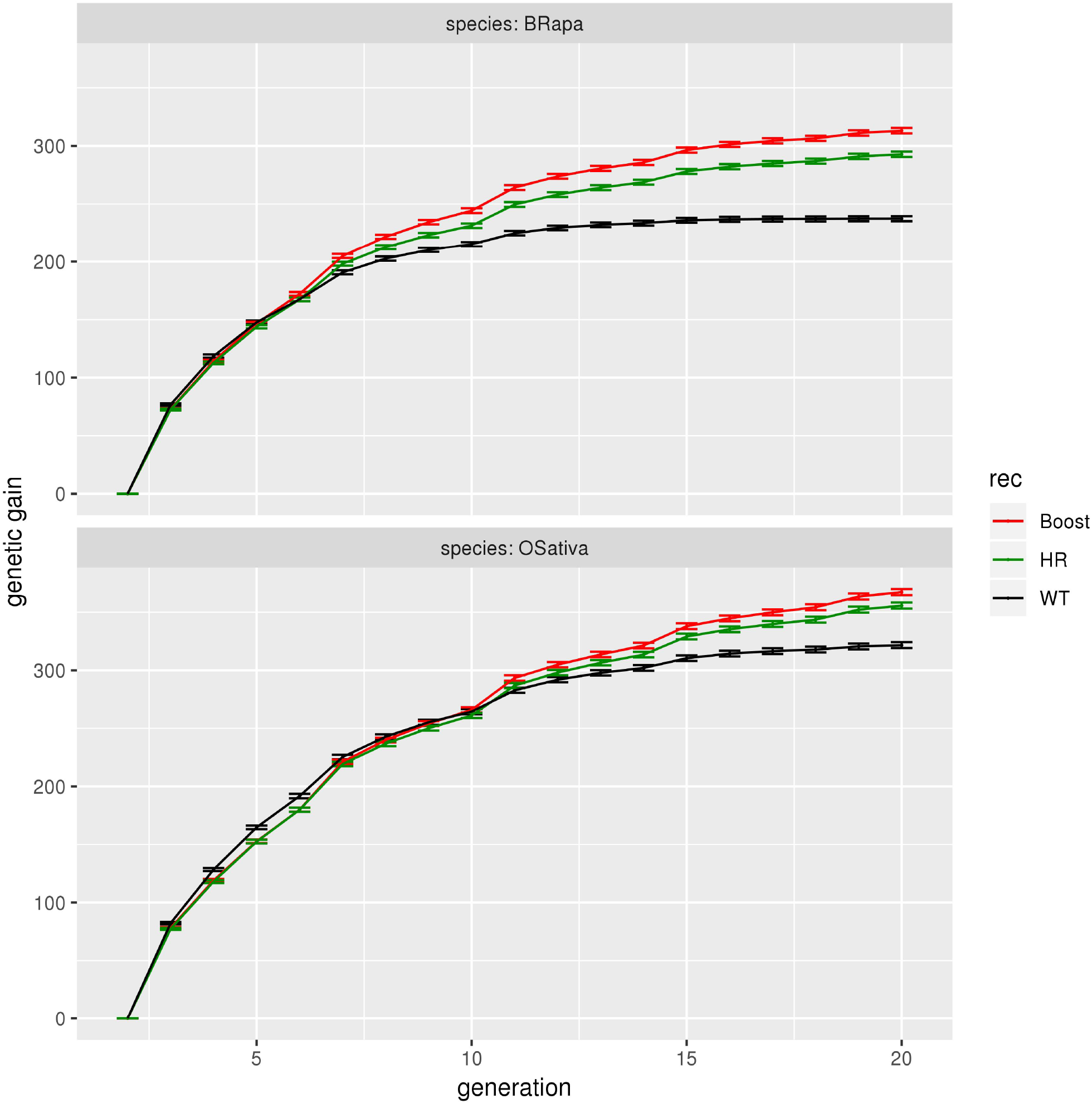
Genetic gain (sum of QTL effects) as a function of the generation number, for *B. rapa* and *O. sativa* under normal (black), HR (green) and boosted (red) recombination, for genomic selection with the marker effects estimated every fourth generation, a heritability of 0.5, selection on heterozygotes with an intensity of 2%, 200 QTLs per chromosome, crossovers formed without interference, no coupling nor repulsion. The gain ratios between increased and normal recombination at g = 20 are as follows. In *B. rapa*, 1.34 for boosted recombination and 1.25 for HR. In *O. sativa*, 1.16 for boosted recombination and 1.12 for HR. The first generation shown is the F2 generation (g = 2). The error bars represent the 95% confidence interval on the mean.

At g = 20, for *B. rapa*, the genetic gain (sum of the QTL effects) is 237 for normal recombination, to be compared to 313 (respectively 293) for boosted (respectively HR) recombination, leading to a gain ratio of 1.34 (respectively 1.25) when using increased recombination. Similarly, at g = 20, for *O. sativa*, the genetic gain is 322 for normal recombination and 367 (respectively 356) for boosted (respectively HR) recombination, leading to gain ratios of 1.16 (respectively 1.12) for that species.

For *B. rapa*, the allelic fixation arises around g = 15 (null genetic variance) under normal recombination, while the genetic gain continues to increase for boosted and HR recombination. Note that the effect of increased recombination on the genetic gain is not immediate. It happens at around g = 7 for *B. rapa* and g = 11 for *O. sativa*. Nevertheless, the effect of increased recombination gets stronger with time. At g = 10, the gain ratio is 1.15 for *B. rapa* under boosted recombination while at g = 20, it is 1.34.

### Re-localized recombination can further increase the gain

Under boosted recombination, there is an increase of the recombination rate in the cold regions, while under HR recombination there is no change in the recombination profile and the recombination is thus mainly increased in the hot regions (Fig1).

Increasing recombination in the cold regions gives us access to more genetic diversity compared to normal recombination. As expected, the effect of increased recombination is stronger under boosted recombination than under HR. For instance, in the case of *B. rapa* (Fig6), the gain ratio is 1.34 for the first *vs* 1.25 for the second. This differential arises even though the gene density in these cold regions is lower than in the rest of the genome (**FigS2**).

### Increased recombination is more efficient when the normal recombination rate is low

Increasing recombination has a stronger effect for *B. rapa* than for *O. sativa*: the gain ratio at g = 20 is 1.34 *vs* 1.16 for boosted recombination and 1.25 *vs* 1.12 for HR (Fig6). The maximum values of the gains being limited by the alleles present in the population, the gain ratios cannot increase indefinitely with increased recombination; for instance the gain ratio between very high and simply high recombination will be close to 1. Instead, the initial level of recombination can be expected to significantly affect the gain ratios and thus it is appropriate to compare those rates in *B. rapa* and *O. sativa*. Since recombination arises both within and across chromosomes, the number of chromosomes (10 for *B. rapa* and 12 for *O. sativa*) will inevitably affect the genetic gain. For instance, in the limit of no intra-chromosomal recombination, it will be advantageous to have more chromosomes. Because of this effect, comparing the genetic gains across these two species is subject to caution. It is possible that the difference in the effect of increased recombination between *B. rapa* and *O. sativa* we see is because of the difference in the genetic lengths, with an average of 80 cM and 275 cM for normal and increased recombination for *B. rapa* and an average of 145 and 475 cM for normal and increased recombination for *O. sativa*. For the same number of QTLs per chromosome, the QTLs in *B. rapa* will be more linked compared to *O. sativa* and so it is plausible that *B. rapa* will respond more strongly to increased recombination.

Given the expectation that increased recombination is most effective if it arises where (normal) recombination rates are lowest, in the following, we choose to concentrate on *B. rapa* under boosted recombination and compare it to normal recombination.

### Heritability does not affect the advantage of increasing recombination

Consider now the influence of heritability on genetic gains in our studies. In the case of phenotypic selection where the sensitivity to heritability should be strongest, it has been previously observed (**Bernardo 2009**, **McClosky and Tanksley 2013**) that heritability plays a rather minor role. In agreement with such a claim, we see in our study of increased recombination that the dependencies on heritability are very small(**FigS4**), with gain ratios of 1.40, 1.38, 1.32, and 1.24 for h^2^ = 1, 0.8, 0.5 and 0.2 at g = 20. The dependencies on heritability are even weaker in the case of genomic selection where the gain ratios are 1.33, 1.37 1.34, and 1.28 at g=20 for those same heritabilities (**FigS5**). The smaller effect of heritability under genomic selection compared to phenotypic selection could have been anticipated because genomic selection estimates the genotypic value and thus attenuates effects of environmental noise.

### Increasing recombination is more efficient with many QTLs

In agreement with what was seen by **McClosky and Tanksley (2013)**, we find that increased recombination is more effective for traits controlled by a lot of QTLs. For instance, there is essentially no effect when there are just 2 QTLs per chromosome, result which can be compared to the gain ratio of 1.34 for 200 QTLs per chromosome in the context of *B. rapa* under boosted recombination at g = 20 (Fig7). This contrast is as expected since the more QTLs per chromosome one has, the more strongly they will be linked and thus the more increased recombination will be effective.

**Figure 7:**
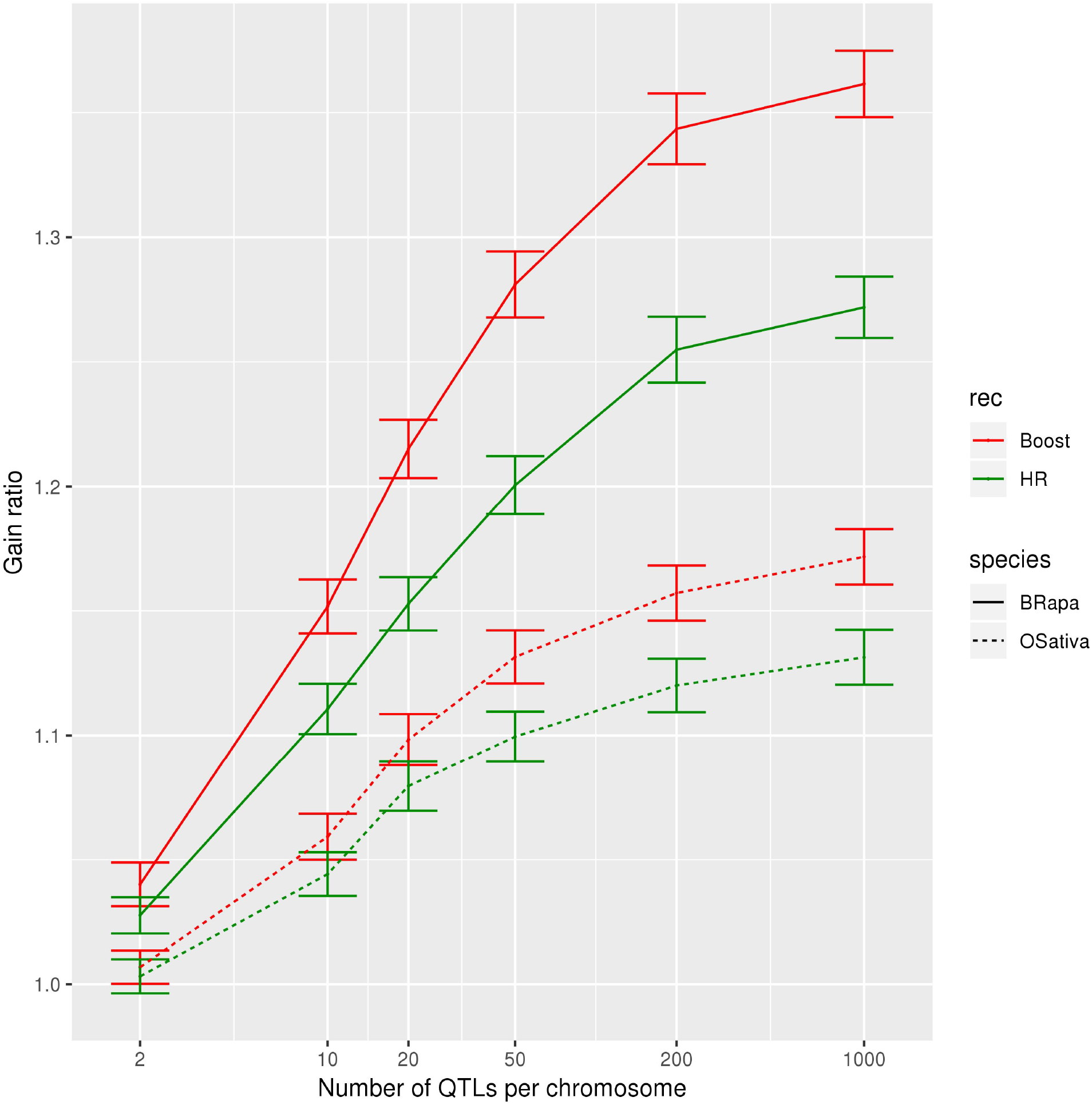
Gain ratio (genetic gain with increased recombination over genetic gain with normal recombination), as a function of the number of QTLs per chromosome, for boosted (red) and HR (green) compared to normal recombination, for *B. rapa* (full lines) and *O. sativa* (dotted lines) for genomic selection with the marker effects estimated every fourth generation, a heritability of 0.5, selection on heterozygotes with an intensity of 2%, crossovers formed without interference, no coupling nor repulsion. The error bars represent 95% confidence interval on the mean.

### Frequent recalibrations improves the benefit of increasing recombination

We also considered the influence of the frequency of recalibrating the genomic selection model. Clearly the best results are expected when the model is recalibrated often, and that is what we see (**FigS6**). But since recalibration requires time and financial resources, the question is to what extent such recalibrations can be spread out in time. In our system, we find that the gain ratio at g=20 between boosted and normal recombination is 1.27 when the model is calibrated *only* in the F2 generation, 1.34 when it is recalibrated once every four generations and 1.35 when recalibrated once every two generations. The difference between these last two cases is not significant, leading us to conclude that there is a regime where one can both benefit from the advantages of genomic selection yet not pay a too high price.

### Increased recombination is more effective under strong selection

Depending on the number of generations used in the selection program, we can afford to select more or less strongly. As up to 20 generations were performed, selection intensities of 2% (5 individuals selected at each generation), 4% (10 selected) and 8% (20 selected) were considered (**FigS7**). In the short-term, higher selection pressures resulted in higher genetic gains (until generation 7 for normal recombination and generation 10 for boosted recombination). However, fixation then happens earlier as a strong selection intensity depletes the diversity faster (fixation at g = 15 under normal recombination with selection at 2%, while fixation is not visible in the other situations). Thus, decreasing the selection intensity and/or increasing recombination delays the fixation. For the last generations, under normal recombination, selecting 4% or 8% of the individuals makes a significant difference while that is not the case under boosted recombination. Hence boosted recombination may be sufficient to conserve a certain level of diversity and thus allow for stronger selection intensity. Increasing recombination is more efficient with a strong selection intensity, the gain ratio between boosted and normal recombination being 1.34 for 2% of selection, 1.30 for 4% and 1.23 for 8% at g = 20.

### Increasing recombination similarly impacts DH and heterozygous selection strategies

All of the previous results were obtained when the selection was done in heterozygous populations (Fn scheme). But in some crops, selection can be performed on doubled haploids (DH) homozygous individuals. Hence, we asked whether our conclusions on the usefulness of increased recombination might extend to the case of selection using DH populations.

Interestingly, for the same number of generations (but half the number of cycles of DH), there is hardly any difference between the two types of populations under selection: at g = 20, the gain ratio is 1.34 for the Fn selection scheme and 1.31 for the DH selection scheme (**FigS8**).

### Increased recombination is beneficial under repulsion but can be detrimental under coupling

So far we supposed a situation without coupling nor repulsion. Since recombination can be expected to be unfavorable in the presence of coupling, it was necessary to quantify the consequences of increased recombination when the alleles were either in coupling or in repulsion (**FigS9**). As expected, under coupling, increasing recombination is detrimental, with a ratio of the gains between boosted and normal recombination of 0.80, while under repulsion, it is beneficial, with a ratio of 1.38, that is more than the ratio of 1.34 under a random situation (neither coupling nor repulsion). Interestingly, we found that the effect of coupling is strong whereas that of repulsion is weak. This may be due to the strong selection intensity (2%). Indeed, under coupling, the selection will result in directly keeping the individuals with large blocks of positive alleles and the gain will have a fast increase. Increasing recombination will break these blocks and lead to a much slower increase of the gain. On the contrary, under repulsion, most of the individuals will have a genotypic value close to zero initially as positive and negative alleles alternate along the chromosomes. Hence, building blocks of positive alleles needs more time, but this is not really different from what happens in the random situation. Nevertheless, in the random situation, some blocks will be in repulsion while others will be in coupling, leading to a higher gain compared to the situation of pure repulsion.

### The benefit of increased recombination remains in the presence of interference

All of the previous results were produced assuming that there was no interference. Having interfering crossovers will lead to fewer events where two adjacent crossovers are close in the same meiosis but it is plausible that this feature has little consequence when considering multiple generations. And this is what transpires from our simulations: increasing recombination still leads to a higher gain (**FigS10**). The effect of boosted recombination is slightly weaker in the presence of interference with a ratio of the gain of 1.31 compared to 1.34 without interference. The interference being lower in the allotriploid AAC compared to the diploid AA of *B. rapa* (lower *nu* parameter of the gamma distribution, **Pelé *et al*. 2017**), the effect of interference on the gain is also lower in the allotriploid. We can also note that the effect of interference on the genetic gain *per se* is small although interference was overestimated (see Material and Method section).

## DISCUSSION

### Increased recombination generally enhances genetic gain across generations

For the range of parameters studied, increased recombination, whether HyperRec or boosted recombination, had a positive effect and led to higher genetic gain than normal recombination. This effect was not observed immediately but took at least three to four generations. However, it got stronger with time for both boosted and HR recombination, which allowed us to conclude that increased recombination slowed down the loss of genetic variance. The early genetic gains (arising during the first generations) may be driven more by random chromosomal assortment than increased numbers of crossovers (**McClosky and Tanksley 2013**). Nevertheless, under recurrent selection, it is more advantageous to increase recombination at the beginning than at the end since recombination acts only when genetic diversity is present. By increasing recombination at the end of program, the effect, if any, would be marginal as most of the diversity would already have been used up.

In our study, the choice of different parameters was important for the size of the effect produced by increased recombination. Among them, the number of QTLs had the highest effect. We found that the genetic gain increased with the number of QTLs, as had already been observed by others (**McClosky and Tanksley 2013**, **Battagin *et al*. 2016**) and such a behavior is expected. Indeed, the more the QTLs one has, the tighter their linkage and the more recombination will be needed to break that linkage. Besides, considering our choice of population size (250 non-independent individuals descending from two homozygous lines), we cannot neglect genetic drift. Hence, the fixation will happen early when there are only a few QTLs, independently of the level of recombination, and increased recombination will have less time to act. It is worth noting that the interaction between genetic drift and selection tends to give an advantage to higher recombination (**Otto and Barton 2001**, **Comeron *et al*. 2008**, **Webster and Hurst 2012**) and thus probably exacerbates the positive effect of increased recombination. Moreover, having an initial population consisting of two parental lines and having the selection start on the F2 population resulted in a high initial linkage disequilibrium, which could also heighten the effect of increased recombination. The strong selection will also heighten the effect of increased recombination as the genetic diversity will be depleted faster and recombination will be needed to keep a certain level of diversity for the selection to act upon. Increasing recombination is thus more interesting in situations where the genetic diversity is limited (initially or due to genetic drift or selection). When considering the others parameters (heritability, selection scheme and types, interference) aside from the number of QTLs, we found that they had only a modest effect on the gain ratio.

### Consequences of increasing recombination in cold regions of the recombination landscape

The highest gain ratio occurred when the regions that are normally cold for recombination were transformed into warm regions. This enhancement to the genetic gain occurred even though the gene density is low in those regions. This result is coherent with the observations made by **Gonen *et al*. (2017)** on the usefulness of shifting recombination hotspots towards the cold regions of the landscapes. Indeed, the absence of recombination in some regions will impede selection there (Hill-Robertson effect, **Hill and Robertson 1966**). By significantly increasing recombination in cold regions, the efficiency of selection in those regions and thus the total effect of selection will increase. It follows that the effect of increased recombination will be most favorable when that increase occurs in cold regions, as observed in our simulations. The major cold regions in many crops are the pericentromeric ones. In fact, these regions can extend to half of a chromosome’s physical length and are almost completely devoid of crossovers (cf. examples by **Sherman and Stack 1995** in tomato, **Anderson *et al*. 2003** in maize and **Choulet *et al*. 2014** in wheat).

Increasing the number of crossovers in these regions are thus predicted by our modeling to lead to significant gains. Note that these crossovers must not be too close to the centromeres if one is to avoid segregation problems and thus loss of fertility (**Talbert and Henikoff 2010**). Such problems might explain the difference of viability and number of seeds per plants observed by **Pelé *et al*. (2017)** in the allotriploids (modified recombination landscape) compared to the diploids. On the other hand, with the HR technology, which almost does not change the recombination landscape and in particular does not put additional crossovers in the cold regions, **Fernandes *et al*. (2018)** did not observe growth or developmental defects in *A. thaliana* but did observe some fertility defects, which were not correlated with the level of increase of recombination. Hence, the fertility problems observed by **Pelé *et al*. (2017)** may not necessarily be due to the increase of recombination near the centromere.

### Feasibility of increased recombination

If increased recombination is to be used in practice, different questions will arise such as when and how should it be applied, and is it worth the cost of doing so? Increasing recombination would incur additional cost, such a producing the mutant for HyperRec or the needed crosses for boosted recombination. As some fertility defects may be expected with increased recombination, one may need to increase the population sizes compared to normal recombination. While HyperRec has been developed in several crops such as pea and tomato (**Mieulet *et al*. 2018**), with varying degrees of increased recombination, boosted recombination has so far only been obtained in *B. rapa*. It is also possible to use other means of increased recombination such as controlling the growth temperature. One may furthermore hope to generate increased recombination through parallel selection for plants with greater number of crossovers as suggested by **McClosky and Tanksley (2013)** but to date there is little experimental evidence that this can be effective. In all cases our modeling predicts that increased recombination has significant effects only when using sufficient generations. Depending on the number of generations wanted, as well as other parameters such as trait architecture, it may not be worthwhile to introduce increased recombination using the kind of methodologies we tested. Instead, it may be more cost effective to resort to methods that have the potential to target recombination to precise locations(**Bernardo 2017**, **Ru and Bernardo 2018**, **Brandariz and Bernardo 2019**). Such approaches are most promising when the trait of interest is controlled by a major cluster of QTLs or when the goal is to produce the proper recombinants in just one generation. However, in this situation of targeted recombination, the precise location of the crossover(s) needs to be determined beforehand. Finally, even though our modeling and simulations indicate how one may best exploit increased recombination (small populations, strong selection intensities, traits controlled by numerous QTLs), it may prove difficult to provide quantitatively reliable predictions because the genetic architecture is typically uncertain and the importance of genetic and environmental interactions that were ignored in our study are generally unknown.

### Consequences for genomic selection programs

**Battagin *et al*. (2016)** wondered about the use of increased recombination for genomic selection and speculated that large datasets would be needed. Indeed, the calibration of genomic selection models is dependent on genomic blocks shared across individuals in the population; increasing recombination reduces the sizes of these blocks. Consequently, the calibration is made more difficult and more importantly the predictive power of the model decreases faster as one accumulates crossovers throughout generations. Nevertheless, our results show a benefit of increased recombination for genomic selection programs even when using a relatively modest training population of 250 individuals. We did observe as anticipated that the accuracy of genomic prediction was reduced under increased recombination (faster break-down of the linkage blocks). But we speculate that under higher recombination the slower loss of genetic variance may compensate the loss of prediction accuracy over generations, justifying the higher genetic gain for boosted and HR recombination than for normal recombination. As **Battagin *et al*. (2016)** expected, increased recombination generated smaller linkage blocks. This led to the estimation of marker effects being closer to the true marker effects compared to normal recombination even though for the generation used for such a calibration the prediction for the genomic value was less good. The greater reliability at the level of the markers resulted in the later generations to a higher genomic prediction accuracy when compared to normal recombination. This was particularly striking in our simulations when no re-estimation was done during selection, the prediction relying entirely on the calibration produced at the F2 generation.

## Acknowledgement

We thank S. Mezmouk, E. Jenczewski, J.C. Montamayor, D. Livingston, C.E. Durel and R. Mercier for constructive comments and discussion.

This work has benefited from a French State grant (LabEx Saclay Plant Sciences-SPS, ANR-10-LABX-0040-SPS), managed by the French National Research Agency under an “Investments for the Future” program (ANR-11-IDEX-0003-02) and KWS, MARS, and Secobra which funded the salary of ET and Ecole Doctorale FIRE - Programme Bettencourt for support.

